# A pathway of NADPH diaphorase positivity between central canal and pial surface at anterior fissure in spinal cord: Supra fissure area with hypothesis configuring from dog, rat, monkey and pigeon

**DOI:** 10.1101/2020.05.02.074450

**Authors:** Yunge Jia, Yinhua Li, Wei Hou, Fuhong Li, Haoran Sun, Xianhui Wu, Xiaoxin Wen, Zicun Wei, Chenxu Rao, Ximeng Xu, Sudirman Fakhruddin Masse, Kuerbanjiang Abulikim, Sheng-fei Xu, Guang-hui Du, Huibing Tan

**Affiliations:** Department of Anatomy, Jinzhou Medical University, Jinzhou, Liaoning 121001, China; Department of Pathology, Heji Hospital Affiliated to Changzhi Medical College, Changzhi, Shanxi, 046011, China; School of Rehabilitation and Sports Medicine, Jinzhou Medical University, 121001, China; Department of Urology, Tongji Hospital, Tongji Medical College, Huazhong University of Science and Technology, Wuhan, Hubei Province, P.R. China; International School, Jinzhou Medical University, Jinzhou, Liaoning 121001, China; Key Laboratory of Neurodegenerative Diseases of Liaoning Province, Jinzhou Medical University, Jinzhou, Liaoning, 121001, China

**Keywords:** NADPH diaphorase, anterior median fissure, central canal, supra fissure area, CSF contacting neurite, spinal cord

## Abstract

The spinal cord is a cylinder structure in the vertebra and thought a simplified with the gray matter and white matter. Rexed lamination for the gray matter and regional sub-division for whiter matter are completely termed to date. Anterior commissure locates between the central canal and the anterior median fissure. However, some experimental data may still confront with new confined anatomical interpretation. By using NADPH diaphorase [N-d] enzyme histology, we found a vertical oriented neuronal pathway between the central canal and the anterior median fissure in the sacral spinal cord of young adult and aged dog. We used a term “supra fissure area” [SFA] to illustrate the region which consisted of the gray commissure and anterior white commissure. The N-d pathway was notably observable in aged animals. The vertical neurites revealed the cerebrospinal fluid [CSF] contacting neurites between the anterior median fissure and the central canal. We further examined the monkey, rat and pigeon in the region for better understanding of the structure and potential function. The neurodegeneration of N-d dystrophy was detected in the [SFA] in the thoracic spinal cord of the aged monkey. N-d positive fibers were detected in anterior fissure of the rat spinal cord. N-d fibrous structures were also detected in the pigeon spinal cord. These results suggested a new pathway of CSF contacting neurons and the neuronal communications about the central canal.

## INTRODUCTION

The spinal cord is composed of the gray matter, white matter and detailed with sub-regional anatomy and physiological function[1]. The development of the spinal cord involves in the folding of the neural plate to form a neural tube[2]. The posterior fissure gradually reduce the central canal in the pig[3]. The anterior median fissure remains until adult[4]. The anterior median fissure of spinal cord is a region of the anterior spinal artery sends perforating branches to supply the central regions of the spinal cord[5]. Both central arteries and central veins circuitously run in the anterior median fissure[6]. One side of dendritic arbors of motoneurons can pass through the anterior commissure travelling to opposite side of anterior funiculus and anterior horn in the spinal cord. The anterior commissure carry projecting fibers from both sides of motoneurons[7]. Most of arborizations of interneurons locate in same side or partially distributed the dorsal commissure[8].

The central canal cellularly organize with ependymal cells and forms ependymal zone as well as sub-ependymal region[9-11]. The nerve plexus occurs the subependymal layer and nerve fibers are located between the ependymal cells[10]. A stem cell’s microenvironment also termed as “niche” is identified around the central canal of the spinal cord[12]. Hamilton et al demonstrate that proliferating ependymal cells are concentrated dorsally in the central canal[9]. The central canal contains the cerebral spinal fluid [CSF]. Selected destroy CSF-contacting neurons may not cause severe disturbances[13]. Syringomyelia is considered in association with the obstruction of CSF flow[14]. This malformation produces some very severe neurological symptoms. Spiller [1909, see review] describes anterior cord syndromed caused by occlusion of the anterior spinal artery or the artery of Adamkiewicz [15]. The regular syndrome affects the anterior 2/3 aspects of the cord. James et al find that the chronic communicating hydrocephalus does not cause enlarge the central canal of the spinal cord and as the alternative pathways for CSF absorption and shows no histological evidence of edema, vacuolation, or tissue destruction [16]. The spinal cord injury adjacent to the ependyma can induce ependymal cells to generate proliferation of ependymal stem/progenitor cells and migrate from the region of the central canal, differentiating into astrocytes [17].

Rexed demonstrate Rexed lamination of the gray matter in the spinal cord by means of regional cytoarchitecture [18]. The white matter is divided into the lateral funiculus, dorsal column, anterior funiculus[4]. The area around the central canal distribute many neuropeptides [19]. Vigh et al report the CSF contacting neurons locate the lateral horn or intermediate zone of the spinal cord and specifically the CSF contacting neurites distribute lateral to the pial surface [20]. Recently, we may find a new neuronal pathway between the central canal to the anterior median fissure by using NADPH diaphorase [N-d]. According to our very preliminary experimental observation, the vertical oriented neurites of N-d positivity were found between the central canal and the anterior median fissure. As matter of fact, spinal cord is a symmetrical structure of side to side pattern for coronal transverse plane, but with the help of the anterior median fissure, perforating branches of the anterior spinal artery supply blood to the central regions of the spinal cord [5]. We figure out a neuroanatomy conception that was hypothesize novel anatomical area termed as supra fissure area based on a pathway of N-d positivity between central canal and pial surface at anterior fissure in spinal cord.

## MATERIALS AND METHODS

### The tissue preparation

The spinal cords of dogs [Canis lupus familiaris, young adult less than 4-year-old, n=5;aged dogs, more than 8-year-old, n=6] of both sexes was legally obtained from Department of Surgery Experimental Animal Facility of Jinzhou Medical University [Jinzhou, China]. The spinal cord of Monkeys [young adult less than 8-year-old, n=4, more than 16-year, n=5] were obtained in the lab of the animal facilities. Male Sprague–Dawley rats [young adult, n=8, 18-month old, n=8] were purchased from Weitonglihua [Beijing, China] and bred in the animal facility of Jinzhou Medical University. The young adult [n = 6] and aged [10-year-old, n = 6] racing pigeons [Columba Livia] were provided and consented by local farmer [Jinzhou, China]. All animal care and experimental procedures met the national guidelines. The experimental protocol was approved by the Ethics Committee on the Use of Animals at Jinzhou Medical University.

The dogs were deeply anesthetized by intravenous injection of sodium pentobarbital [overdosage of 60 mg/kg body weight]. Briefly, the dogs were trans-cardially perfused through the aorta with normal saline, followed by 4% paraformaldehyde in 0.1 M phosphate buffer [PB; pH 7.4]. The tissues of the spinal cord were rapidly obtained and immersed in the same fixative as in the perfusion for at least 6 hrs. at 4°C, and transferred to 30% sucrose in 0.1 M PB [pH 7.4]. Frozen sections were cut transversely or horizontally at 40 µm on a cryostat [Leica, German]. Similarly, rats and pigeons were deeply anesthetized with intraperitoneal injection of sodium pentobarbital [50 mg/kg]. The spinal cords were obtained after antheses. The tissues were processed with coronal and sagittal sectioning respectively.

### NADPH diaphorase histochemistry

N-d enzyme-histology was performed in free-floating method. In this procedure, sections were incubated for 5 min in 100 mM sodium phosphate buffer PBS, pH 7.4] followed by incubation in phosphate buffer [PB, pH 7.4] with 1 mM beta-NADPH [Sigma, USA], 0.5 mM nitrotetrazolium blue [NBT, Sigma, USA] and 0.3% Triton X-100 at 37°C for up to 2-4h. Sections were rinsed with PB, distilled water, dehydrated in a graded ethanol series, and were coverslipped with mounting medium.

### Quantitative microscopy and Statistical analysis

Sections were observed under the light microscope [Olympus BX53 microscope, Japan]. Images were captured with a DP80 camera. Sections from all spinal cord levels in each animal were quantitated using Olympus image analysis software [Cellsens Standard, Olympus].

## RESULTS

We used N-d staining to identify subpopulations of neurons in the spinal cord[21]. We also found some positive neurons were detected in the anterior horn[22]. Recently, neuronal circuit of N-d positivity are considered as functional structures in both normal and aging organizations[23-26]. The present study aimed to specify hypothetic observation: supra fissure area between the central canal and the anterior median fissure the spinal cord. In order to define the morphological location, we cited two illustrations to show the supra fissure area [SFA] [Figure 1.] from Clarke[27] and Cajal [28]. *Supra fissure* in the term indicated orientation because the most transverse sections of spinal cord are showed dorsal to ventral oriented pattern. The space of the SFA was set with a top boundary reached to the central canal, a lower boundary set to the top and around the anterior median fissure and two side boundaries between the anterior [ventral] horns of gray matter. The Figure 1 shows many decussated fibers in the SFA. We found some vertical N-d fibers oriented between the central canal and the anterior median fissure.

**Figure 1.**
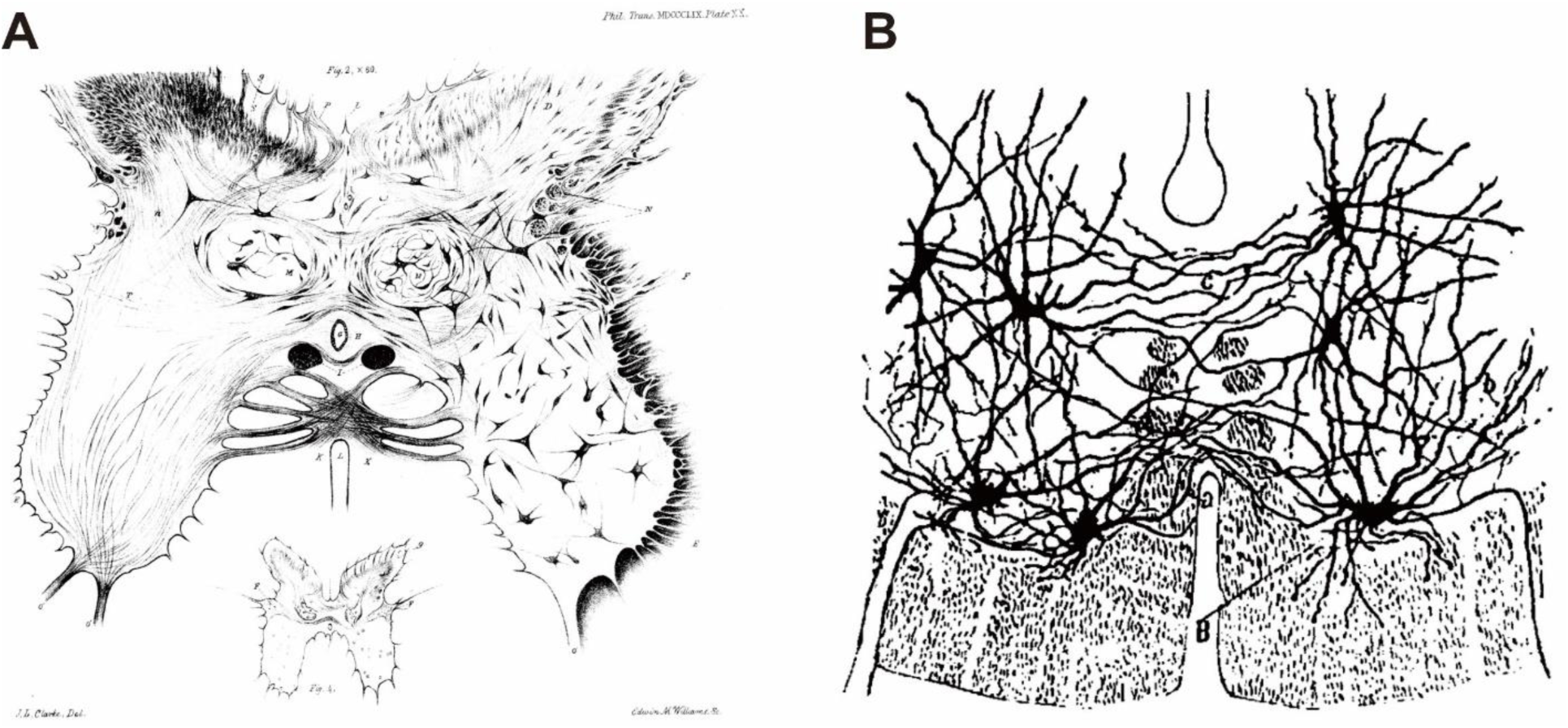
Adapt two illustrations to demonstrate of the region between the central canal and the anterior median fissure [AMF]. A showed the drawing of the sacral spinal cord in ox from Clarke[27]. Note the region [supra fissure area] between the AMF and the central canal. B: Drawing adapted from Cajal [28] of commissural nerve cells in the medial part of the ventral horn[18]. In this illustration, A indicated drorsomedial nerve cells with neurites crossing the anterior commissure, while B indicated ventromedial neurons sending its neurites into the ventral root.

We first stated our hypothesis from the observation of the spinal cord of adult dog. Figure 2 showed N-d positivity in the caudal sacral spinal cord in young adult dog. Dendrites are the input site of neural signals into neuronal somas. We noted that some thick dendrites emerged proximal to the neuronal cell bodies [Figure 2 B and C] in the coronal section of the caudal sacral spinal cord. The segments of thick neurites were detected in Figure 2 B in both the dorsal commissural nucleus and the supra fissure area, while thin fiber in the supra fissure area. The vertical oriented neurites both thick and thin fibers occurred in the supra fissure area [open arrowhead in B and double thin arrows in E]. An interesting claw-like neurites in the central canal [Figure 2 G, open arrow]. It was definitely a cerebrospinal fluid [CSF] contacting terminals in the central canal. Next, we examined the same segment of the sacral spinal cord in aged dog. As the previous investigation, the ANBs and megaloneurites were detected in the sacral spinal cord in aged dog [24, 29]. The caudal sacral spinal cord occurred relative a smaller number of the megaloneurites than the rostral sacral. However, the neurites in the supra fissure area and intra-central canal N-d positivity were still quite visible in the caudal sacral segment [Figure 3]. The megaloneurites were detected in the lateral collateral pathway, area around the central canal and the anterior median fissure [Figure 3 B]. It was also a CSF contacting neurite to external CSF. It may be the evidence for non-synaptic communication[30] or the “sink” of the central canal[31]. In the same photo image, thin fiber also was visualized along the other side of the anterior median fissure. More dorsal to ventral oriented neurites revealed a neurol circuit pathway between the central canal and the anterior median fissure in the Figure 4. The aging-related transverse of the megaloneurites occurred in the dorsal lateral funiculus, which were consistent our previous discovery[22] [Figure 4 B,D and H]. The central canal N-d positivity was also detected. Some neurites located intra-ependymal cells. At least three neurites/ megaloneurites showed in three sections located between the central canal and anterior median fissure. It was a typical illustration to characteristic supra fissure area. Another two examples of cerebrospinal fluid contacting neurites between the central canal and the anterior median fissure in Figure 5 were further verified N-d pathway between the central canal and the anterior median fissure at the caudal spinal cord. The relevant neurites extended along the pial surface of the anterior median fissure and form plexus. Similar intra-ependymal neurites were also detected. The neurites in supra fissure area could reach both the interior CSF in the central canal and exterior CSF surround spinal cord.

**Figure 2.**
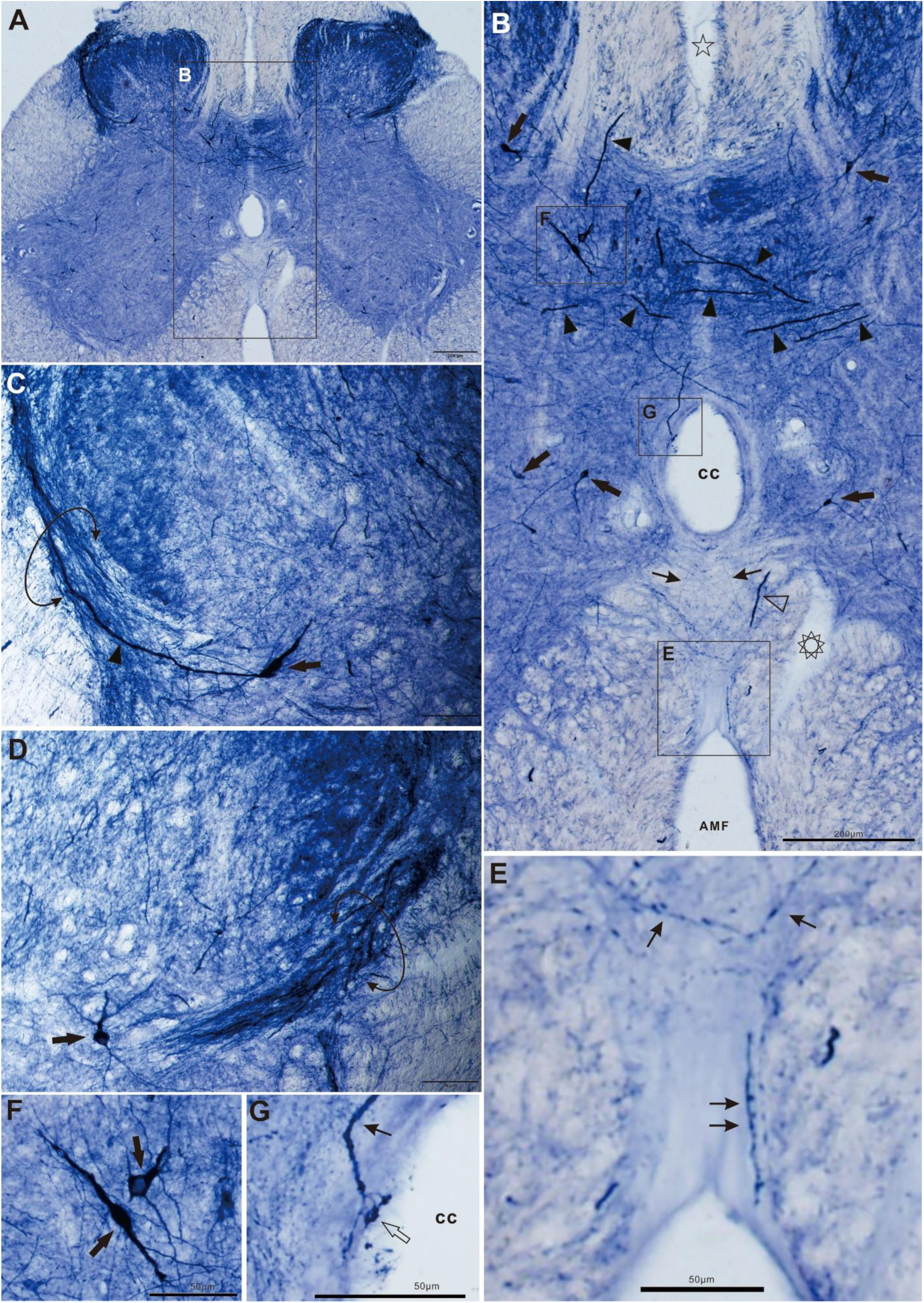
N-d positivity in the sacral spinal cord in young adult dog. A: the coronal section of the sacral spinal cord. B: Magnification from A. Star showed the perforating branch of the posterior vessels surround by N-d positive fibers. Complex polygon indicated the perforating branch of the anterior vessels. Arrow indicated neuron, arrowhead indicated thick neurite, thin arrow indicated thin fiber in the supra fissure area. cc indicated the central canal. AMF indicated anterior median fissure. C and D showed the lateral collateral pathway [circle arrow] in the dorsal horn. B, F and G magnified of B. Double arrow in E showed a vertical neurite. G showed claw-like neurites in the central canal [open arrow]. Bar in A and B =200μm, C-G =50μm.

**Figure 3.**
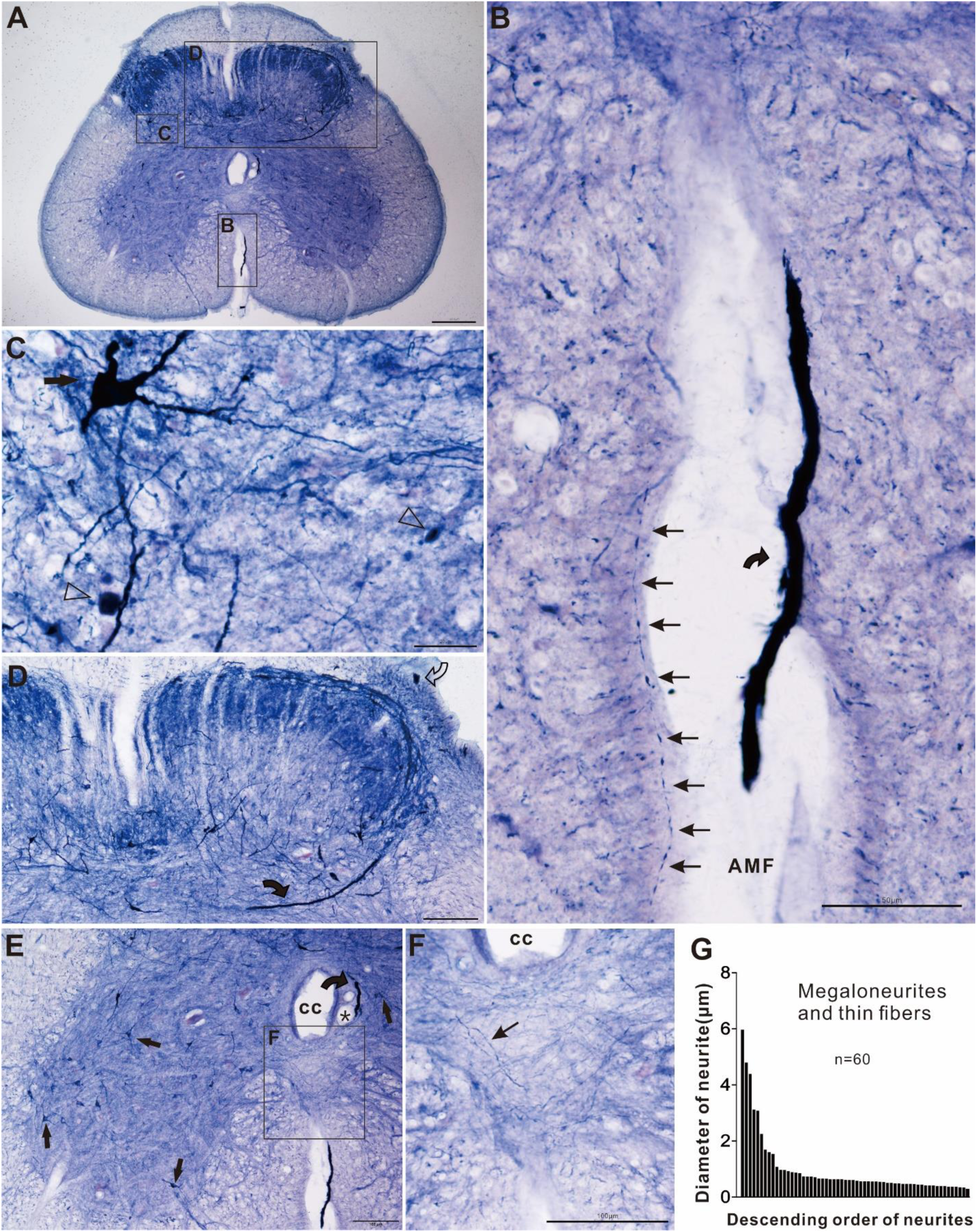
Megaloneurites detected in the anterior fissure in aged dog. A showed the coronal section of the sacral spinal cord. B magnified from A showed the megaloneurite [curved arrow] in the anterior median fissure [AMF]. The linear arranged thin arrow indicated subpial varicose thin fiber. C showed intermediolateral nucleus. Arrow indicated neuron. Open arrowhead indicated ANB. D showed the segment of megaloneurite [curved arrow] in Lissauer’s tract and megaloneurite [curved arrow] in the lateral collateral pathway. E showed ventral horn and central canal [cc] as well as the AMF. Curved arrow indicated megaloneurite. Asterisk indicated blood vessel. F indicated the supra fissure area. Thin arrow indicated an example of N-d fiber. Histogram G showed diameter of the neurites from the D. Bar in A =200μm, in B, D, E and F =100μm, bar in C=20μm.

**Figure 4.**
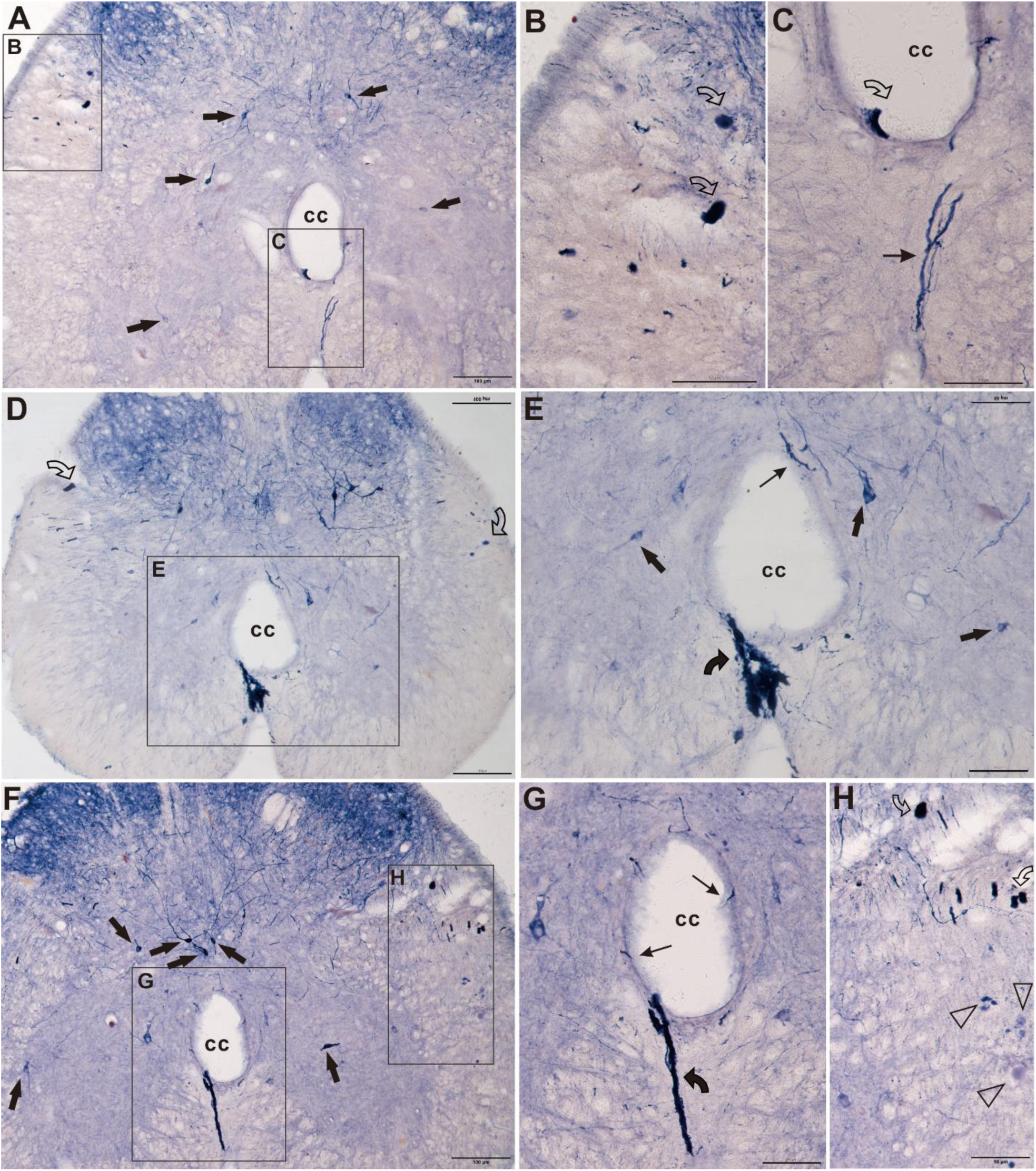
N-d pathway showed between the central canal [cc] and the anterior median fissure [AMF] in aged dog. A, D and F showed caudal segment of the spinal cord. Arrow indicated neuron. B, C, G and H showed similar pattern for transverse megaloneurites [open curve arrow]. C showed thick neurite [thin arrow]. E and G showed megaloneurite [curve arrow]. Bar in A, D and F =100μm, the other = 50μm.

**Figure 5.**
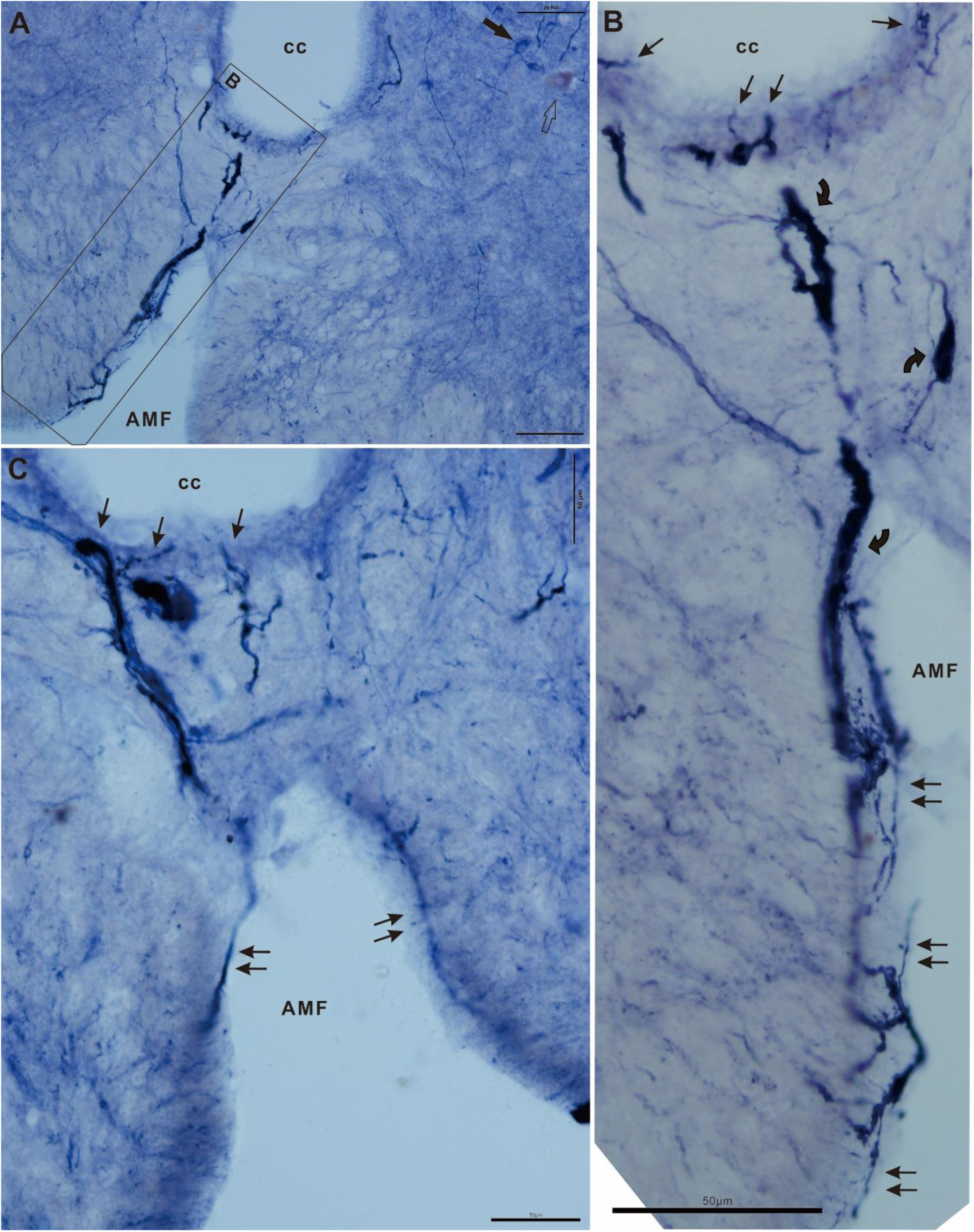
Another two examples to demonstrate N-d pathway between the central canal [cc] and the anterior median fissure [AMF] at the caudal spinal cord. A showed the N-d neurites in the region between the cc and AMF and intra-ependymal cells as well as the sub-ependymal layer. Arrow indicated neuron. Open arrow indicated very weak stained somatic neuron. Note that the neurites in the rectangle formed plexus in pial surface. B magnified from A. Thin arrow indicated neurites between the ependymal cells. Curved arrowed indicated megaloneurite. Double arrow indicated fiber in the pial surface. B showed similar distribution pattern of neurites. Bar = 50μm.

After examining the sacral spinal cord of dog, the spinal cords of monkey were examined. We found that dorsal to ventral oriented fibers and neuronal soma in the supra fissure area in the transverse section of the sacral spinal cord [Figure 6 D and E] and intra-ependymal neurites in the central canal as well as CSF-contacting fiber in subpial or surface along the anterior median fissure of spinal cord in adult monkey [data not showed here]. In thoracic spinal cord of aged monkey, some neuronal dystrophic neurites or spheroids occurred in the supra fissure area. The neurodegeneration in the supra fissure area implicated dysfunction of autonomous nervous system in aging.

**Figure 6.**
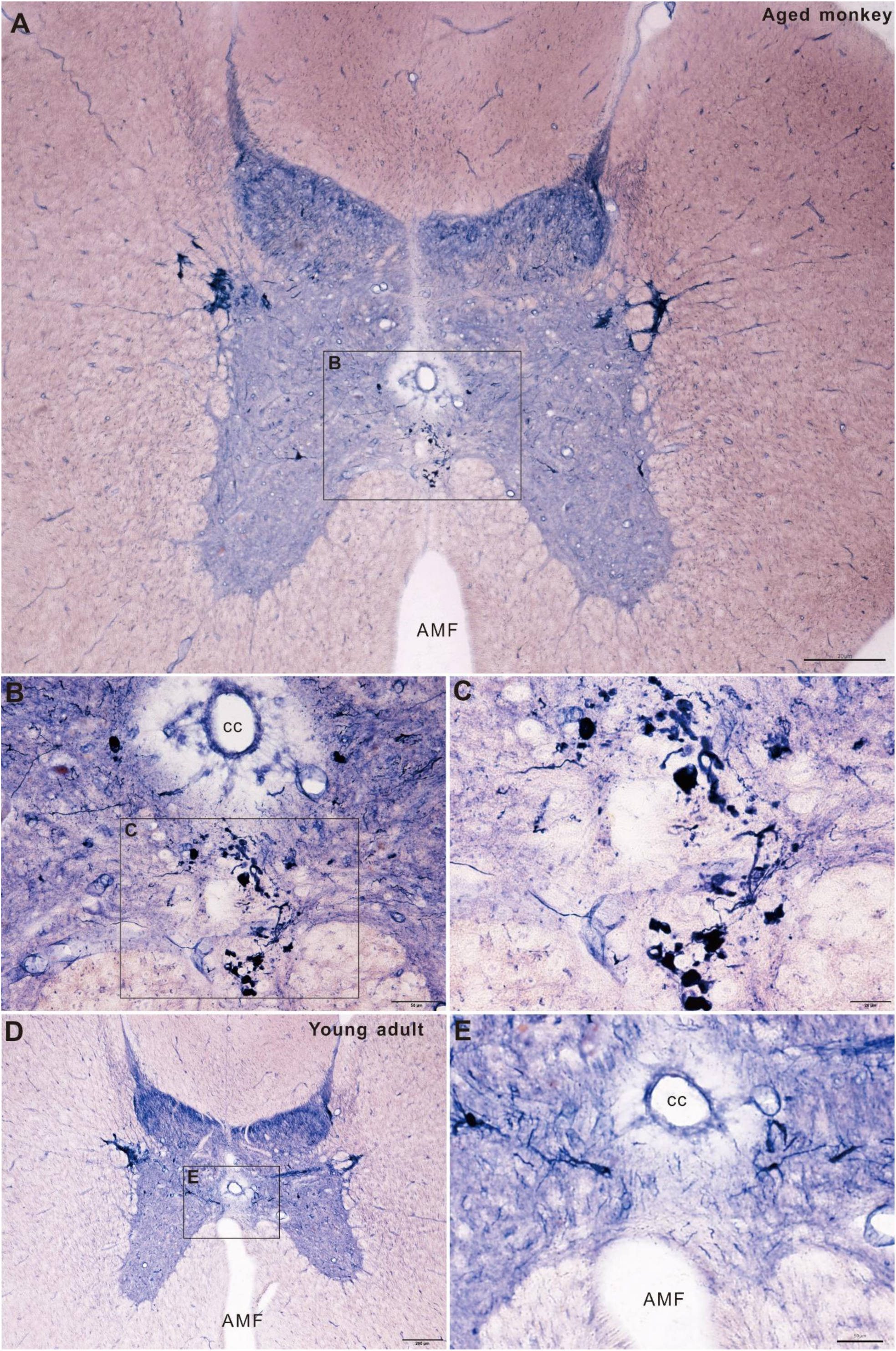
Detected ANB in the supra fissure area the thoracic in the spinal cord of aged monkey. A-C showed ANB in the supra fissure area. B and C magnified from A. D and E to show the same region of the supra fissure area in young adult money. E magnified from D. Bar A and D =200μm, B and E =50μm, and C=20μm.

Rat is commonly used experimental animal. Normally, the vertical oriented fibers and intra-ependymal neurites were also detected between the central canal and the anterior median fissure in young adult rats. We do not find the megaloneurites in aged rat. The aging-related N-d bodies were found in the spinal cord as the previous study[32]. We still focused on examining the supra fissure area in rat. In sagittal section of spinal cord in aged rat, vertical oriented fibers were detected through the anterior spinal artery at the thoracic segment [Figure 7].

**Figure 7.**
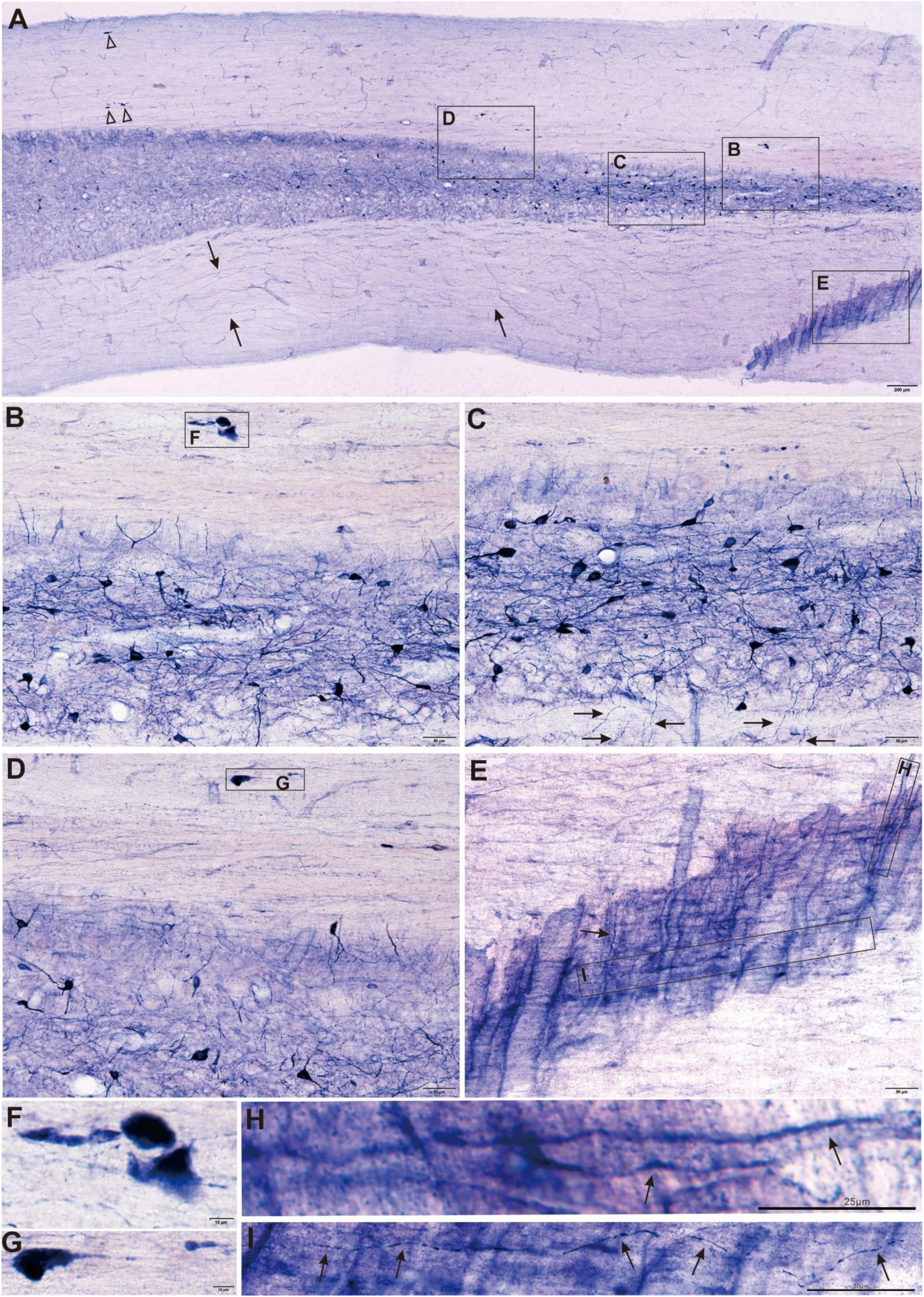
Sagittal section through the partial anterior spinal artery at the thoracic segment of aged rat. A: Region E showed the anterior spinal artery. Open arrowhead indicated ANB in the dorsal column. Arrow indicated example of the longitudinal fibers some of which extended around 2/5 of the section. B, C and E magnified from A. Arrow in C indicated vertical fiber toward to anterior fissure. The different cellular and fiber intensity detected between the ventral and dorsal gray matter [A-C]. F and G magnified from B and D respectively, and indicated ANBs. E showed the anterior spinal artery. H showed the vertical N-d fiber [arrow] nearby the anterior spinal artery. I showed a longitudinal fiber [arrow]. Bar in A= 200μm, B-G and I =50μm, H= 25μm.

The anterior median fissure is a common feature of the spinal cord in most mammals. For pigeon, a unique splitting of the dorsal column to form the rhomboid sinus reveals in the lumbosacral enlargement[33]. The distance between the central canal and the anterior fissure was short in lumber segment of pigeon, while the depth of the anterior fissure was relatively superficial to ventral boundary of the lumber spinal cord. The N-d fibrous positivity revealed between the central canal and the anterior median fissure [Figure 8 A-C and D]. We noted a uvula shaped structure at the bottom of the supra fissure area [Figure 8 C]. The intuitional reasoning for the uvula structure was something like a rudder on a boat. There was no uvula structure showed in the Figure 8 E. Maybe it caused by mounting. Figure 9 showed the supra fissure area in the cervical spinal cord of adult pigeon. Figure 9 A-D showed neurons and fibers in the supra fissure area. Some dendrites of neurons tracing to the supra fissure area, which were different to the bridging connections between the central canal and anterior median fissure. Figure 9 E-H showed another example of the supra fissure area. A high-density N-d reaction showed at the dorsal end of the anterior mediate fissure.

**Figure 8.**
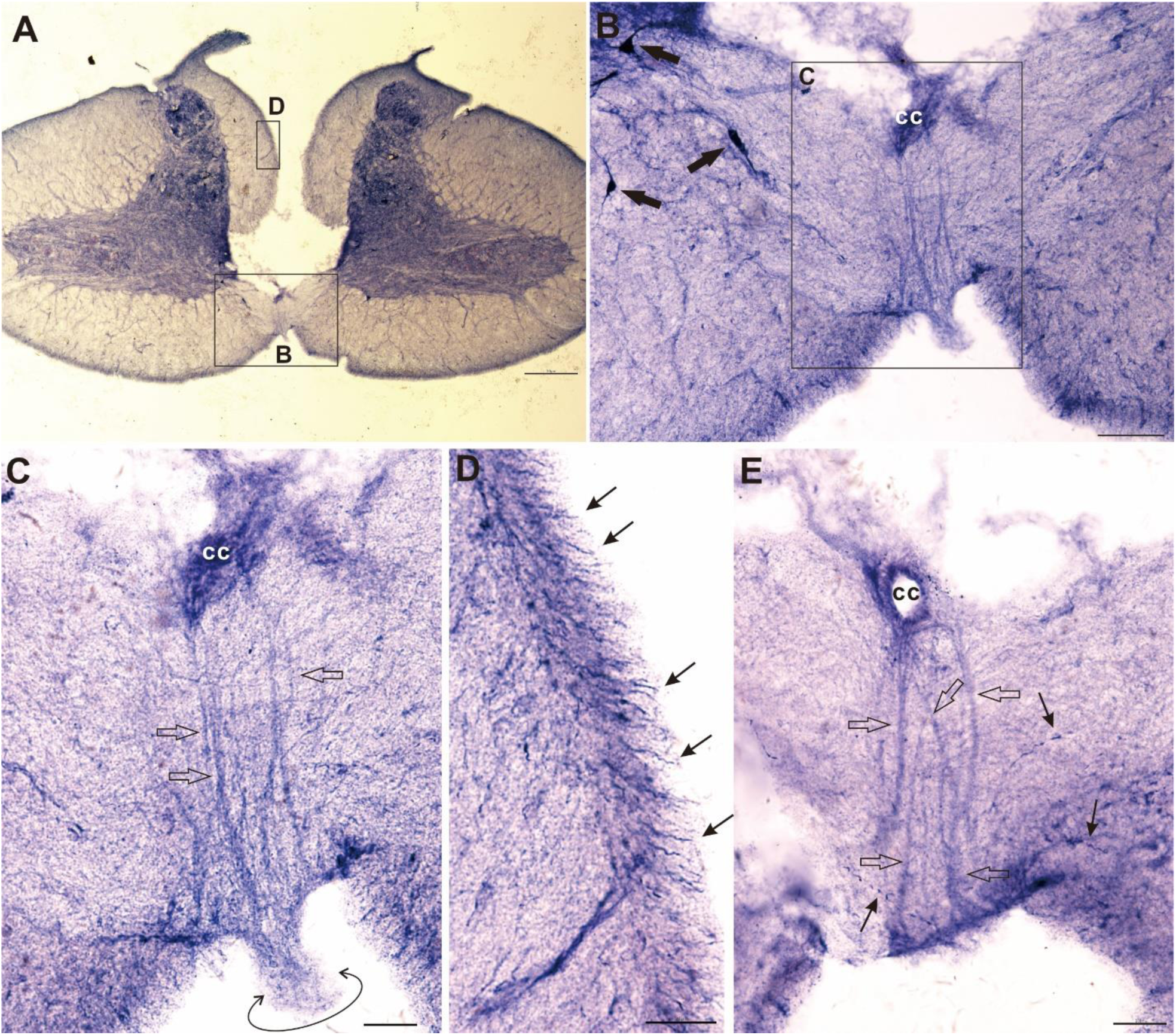
N-d fibrous activities detected between the central canal [cc] and the anterior median fissure [AMF] at the lumber spinal cord of adult pigeon. A-C showed lightly stained thick fibers. Arrow indicated neurons. Open arrow indicated N-d thick fibers between cc and AMF. The double circle arrow indicated a uvula-like structure in the AMF. D magnified from A which showed pial fibers [arrow]. E was another example for N-d thick fibers between cc and AMF. Bar in A =200μm, B=50μm, C-E=20μm.

**Figure 9.**
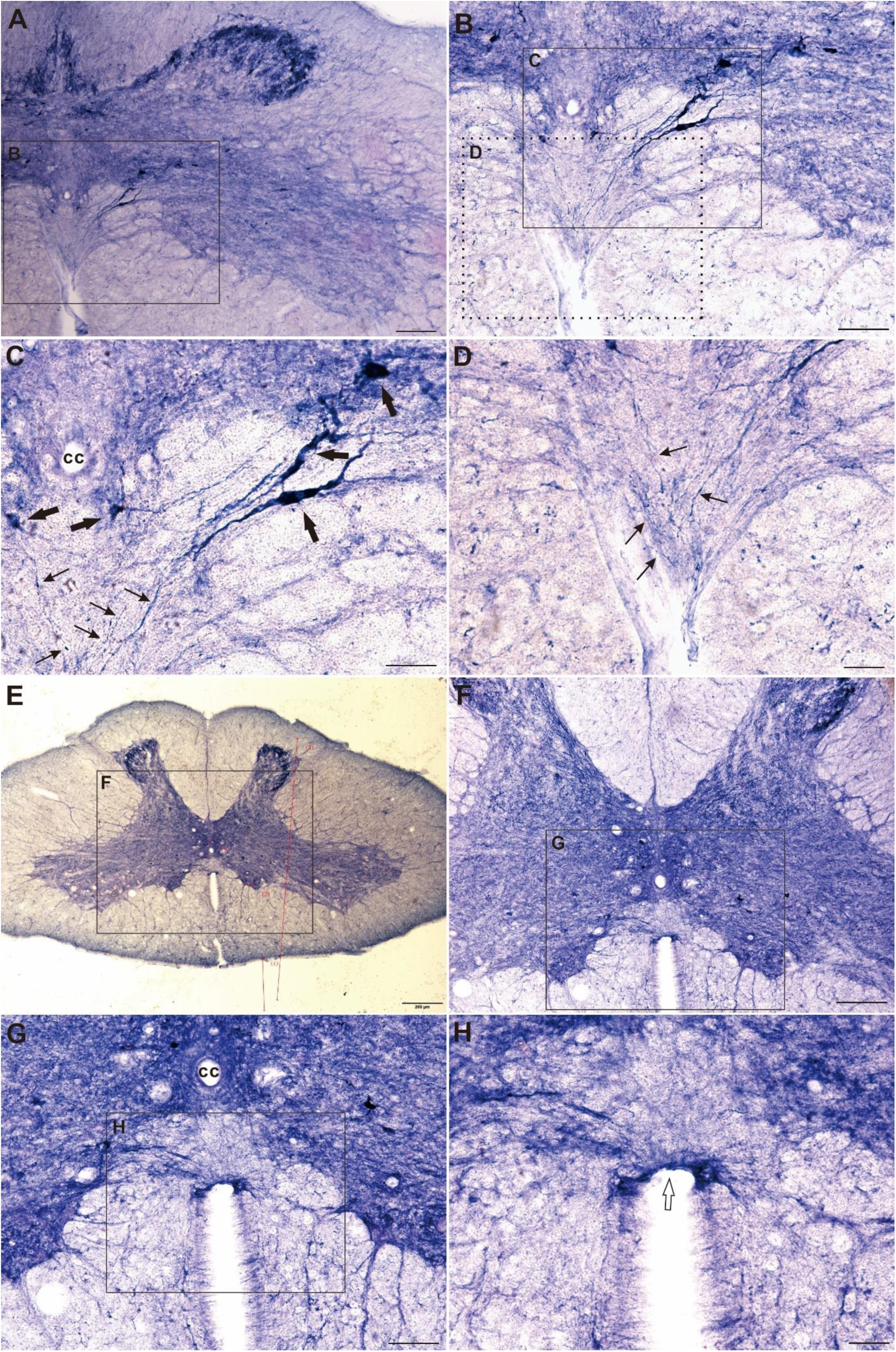
Supra fissure area in the cervical spinal cord of aged pigeon. A-D showed neurons and fibers in the supra fissure area. Arrow indicated neuron. Thin arrow indicated fiber. cc: central canal. E-H showed another example of the supra fissure area. Open arrow indicated a high-density N-d reaction at the dorsal end of the anterior mediate fissure. Bar in A and E=200μm, B and F=100μm, C and G=50μm, D and H=20μm.

## DISCUSSION

To identification the specialized neurons and fibers with N-d staining, we found that some N-d neurites distributed between the central canal and the anterior median fissure. We further demonstrated a hypothesis of unrecognized region named as the supra fissure area. The primitive observation based on the N-d staining of the sacral spinal cord of aged dog because it showed megaloneurites and thick neurites in the supra fissure area. In general, the previous illustrations of Clarke[27] and Cajal [28] do not show any “intrinsic” relation vertical oriented neurites between the AMF and the central canal. N-d staining in young adult dog may minorly illustrate the N-d positivity to indicate a neuronal pathway between the central canal and anterior median fissure. It still is uncertain to make a conclusion of neuronal pathway between the AMF and the central canal. The aging change of megaloneurites in aged dog coincided the fact that a pathway of N-d staining between the central canal and anterior median fissure. For N-d staining in pigeon, the uvula shaped structure in the lumber segments may be still a counterpart as a sense organ of equilibrium[34, 35].

CSF flow through the central canal of the spinal cord[36]. Milhorat et al think that internal CSF flows in a rostral direction through the central canal of the spinal cord in rats[37]. In computerized 3-D study, caudal spinal cord forms *canal duplication, a terminal ventricle and openings from the canal lumen into the subarachnoid space*[31]. Some investigators *suggest the possibility of a functionally important fluid communication in the caudal spinal cord which may have a sink function* [31, 38-40].

The neurites of CSF-contacting neurons showed horizontal orientation in the spinal cord in frog embryos [41]. The processes of tanycytes in the central canal extend dorsal and ventral orientation [42]. Hypothalamic tanycytes in the ventral portion of the third ventricle were NADPH-d positive the leptin systemic administration[43]. Visceral and somatic afferent convergence onto neurons near the central canal in the sacral spinal cord of the cat, some neurons in the dorsal commissural nucleus send neurites around the central canal projecting to the anterior commissure [44]. Intraventricular injection of horseradish peroxidase, horseradish peroxidase positivity distributed in the rat lumbosacral spinal cord[45]. Central canal may connect with blood vessels[45].

The pia-arachnoid associated tissue and cellular component play important in the development of both the posterior median septum[sulcus] and the anterior median fissure[46]. The pia-arachnoid forms a prolongated connection for the spinal CSF with the venous blood drainage which may communicate either through the anterior and posterior spinal veins or through the central veins. Our finding and hypothesis concerned that the N-d pathway between the central canal and the anterior median fissure. Besides the N-d pathway, it could exist other neurites. We thought that the anterior median fissure made a shortcut or alternate route to provide convenient and efficient arrangement between internal CSF in the central canal and the external CSF surrounding the spinal cord. The anterior median fissure definitely works as harbor of blood supplier and docking the main artery and vain. The N-d neurons and fibers also distribute the ventral of the human spinal cord and the region above of the anterior median fissure[47].

It is common to think of white matter as the site of descending and ascending pathway composed of axonal bundles in the spinal cord. However, the white matter forms a certain topographic function unit. The posterior funiculus or dorsal column distributes not only dorsal column funiculus [gracile funiculus and cuneatus funiculus] but also neuronal cells, which function as integration of neuronal circuit [48]. In our recent investigation, the N-d CSF contacting neurons and plexus as well as subpial neurons in the median dorsal funiculus configurate a CSF detecting and sensation structure[25]. What function could be for the N-d pathway between the central canal and the anterior fissure? Summary from the previous investigations[20, 49], CSF contacting neurons may response for the pressure, pH value, osmolarity and chemicals as well as neurotransmitters in CSF. hydrocephalus and canal occlusion[50]. Most of the CSF contacting neurons as tanycytes send their apical neurites toward ependymal surface[51]. We noted that some vasoactive intestinal peptide [VIP] positive neurons distribute among the ependymal cells or sub-ependymal cells around the central canal[52, 53]. The VIP-ergic fibers also distribute the anterior fissure[53]. The distribution of the N-d fibers kept consistent with the VIP-ergic fibers distributed in the around the central canal and the anterior fissure. In our previous study, we also find that the VIP-ergic fibers distribute around the central canal[29]. Meanwhile, N-d megaloneurites and some fibers co-localize with VIP[22, 54]. Some VIP positive fibers and megaloneurites distribute toward to the anterior commissure[29]. In the middle of the central canal, there is a thread-like structure named Reissner’s fiber extending from the subcommissural organ to the entire central canal in the spinal cord[55]. N-d is supposed to identical to the nitric oxide synthase [NOS][56, 57]. However, N-d positive staining for the subcommissural organ is negative to NOS immunoreactivity[58]. The subcommissural organ is important to form the Reissner’s fiber in the central canal. We thought that the central canal should be considered as a crucial organ for CSF circulation and motion control[25].

In summary, in order to initiate discussion of the speculated idea, we termed the region of the supra fissure area. the supra fissure area between the central canal and the anterior median fissure was examined in dog, monkey, rat and pigeon to identify features that distinguished them regionally. The most notable were the megaloneurites between the central canal and the anterior median fissure in aged dog, degenerated spheroid in the monkey and thick fibers in pigeon. The results revealed that the communications of internal CSF in the central canal with external CSF in the space surround the spinal cord.

## ACKNOWLEDGMENTS

This work was supported by grants from National Natural Science Foundation of China [81471286] and Research Start-Up Grant for New Science Faculty of Jinzhou Medical University [173514017].

## References

[1] M.K. Ganapathy, P. Tadi, Neuroanatomy, Spinal Cord Morphology, StatPearls [Internet], StatPearls Publishing 2019.

[2] K.W.S. Ashwell, Chapter 2 - Development of the Spinal Cord, in: C. Watson, G. Paxinos, G. Kayalioglu (Eds.) The Spinal Cord, Academic Press, San Diego, 2009, pp. 8–16.

[3] W. Barnes, On the Development of the Posterior Fissure of the Spinal Cord, and the Reduction of the Central Canal, in the Pig, Proceedings of the American Academy of Arts and Sciences, JSTOR, 1883, pp. 97–110.

[4] N. Goto, N. Otsuka, Development and anatomy of the spinal cord, Neuropathology, 17 (1997) 25–31.

[5] G. Romanes, The arterial blood supply of the human spinal cord, Spinal Cord, 2 (1965) 199–207.

[6] Y. Naka, T. Itakura, K. Nakai, K. Nakakita, H. Imai, T. Okuno, I. Kamei, N. Komai, Microangioarchitecture of the feline spinal cord, 66 (1987) 447.

[7] A.R. Light, C.B. Metz, The morphology of the spinal cord efferent and afferent neurons contributing to the ventral roots of the cat, Journal of Comparative Neurology, 179 (1978) 501–515.

[8] M. Matsushita, Some aspects of the interneuronal connections in cat’s spinal gray matter, Journal of Comparative Neurology, 136 (1969) 57–79.

[9] L. Hamilton, M. Truong, M. Bednarczyk, A. Aumont, K. Fernandes, Cellular organization of the central canal ependymal zone, a niche of latent neural stem cells in the adult mammalian spinal cord, Neuroscience, 164 (2009) 1044–1056.

[10] K. Nandy, G. Bourne, Histochemical studies on the ependyma lining the central canal of the spinal cord in the rat with a note on its functional significance, Cells Tissues Organs, 60 (1965) 539–550.

[11] J.E. Bruni, K. Reddy, Ependyma of the central canal of the rat spinal cord: a light and transmission electron microscopic study, Journal of anatomy, 152 (1987) 55–70.

[12] Y.W. Zichun Wei, Ximeng Xu,Wei Wu, Chenxu Rao, Huibing Tan, Poterntial of spinal cord ependymal cell regeneration and other related cell targeted therapuetic strategies, Chin J Neurodrauma Surg, 1 (2015) 50.

[13] S.Y. Song, L.C. Zhang, The Establishment of a CSF-Contacting Nucleus “Knockout” Model Animal, Front Neuroanat, 12 (2018) 22.

[14] K.H. Støverud, M. Alnæs, H.P. Langtangen, V. Haughton, K.-A. Mardal, Poro-elastic modeling of Syringomyelia–a systematic study of the effects of pia mater, central canal, median fissure, white and gray matter on pressure wave propagation and fluid movement within the cervical spinal cord, Computer methods in biomechanics and biomedical engineering, 19 (2016) 686–698.

[15] E. Diaz, H. Morales, Spinal cord anatomy and clinical syndromes, Seminars in Ultrasound, CT and MRI, Elsevier, 2016, pp. 360–371.

[16] A.E. James, W.J. Flor, G.R. Novak, E.-P. Strecker, B. Burns, Evaluation of the central canal of the spinal cord in experimentally induced hydrocephalus, 48 (1978) 970.

[17] A.J. Mothe, C.H. Tator, Proliferation, migration, and differentiation of endogenous ependymal region stem/progenitor cells following minimal spinal cord injury in the adult rat, Neuroscience, 131 (2005) 177–187.

[18] B. Rexed, The cytoarchitectonic organization of the spinal cord in the cat, The Journal of comparative neurology, 96 (1952) 415–495.

[19] S.J. Gibson, J.M. Polak, S.R. Bloom, P.D. Wall, The distribution of nine peptides in rat spinal cord with special emphasis on the substantia gelatinosa and on the area around the central canal (lamina X), The Journal of comparative neurology, 201 (1981) 65–79.

[20] B. Vigh, I. Vigh-Teichmann, B. Aros, Special dendritic and axonal endings formed by the cerebrospinal fluid contacting neurons of the spinal cord, Cell and Tissue Research, 183 (1977) 541–552.

[21] C.R. Anderson, NADPH diaphorase-positive neurons in the rat spinal cord include a subpopulation of autonomic preganglionic neurons, Neuroscience letters, 139 (1992) 280–284.

[22] Y. Li, Y. Jia, W. Hou, Z. Wei, X. Wen, Y. Tian, W. Zhang, L. Bai, A. Guo, G. Du, H. Tan, De novo aging-related megaloneurites: alteration of NADPH diaphorase positivity in the sacral spinal cord of the aged dog, bioRxiv, DOI 10.1101/483990(2019) 483990.

[23] W. Hou, Y. Jia, Y. Li, Z. Wei, X. Wen, C. Rao, X. Xu, F. Li, X. Wu, H. Sun, H. Li, Y. Huang, J. Sun, G. Shu, X. Wang, T. Zhang, G. Shi, A. Guo, S. Xu, G. Du, H. Tan, NADPH diaphorase neuronal dystrophy in gracile nucleus, cuneatus nucleus and spinal trigeminal nucleus in aged rat, bioRxiv, DOI 10.1101/2019.12.21.885988(2019) 2019.2012.2021.885988.

[24] Y. Li, Y. Jia, W. Hou, H. Tan, Novel Morphological Alteration and Neural Pathway of NADPH Diaphorase Positivity in the Spinal Cord with Disc Herniation of Aged Dog: A Case Report, DOI (2020).

[25] Y. Li, W. Hou, Y. Jia, X. Wen, C. Rao, X. Xu, Z. Wei, L. Bai, H. Tan, Pial surface CSF-contacting texture, subpial and funicular plexus in the thoracic spinal cord in monkey: NADPH diaphorase histological configuration, bioRxiv, DOI 10.1101/2020.01.30.927509(2020) 2020.2001.2030.927509.

[26] Y. Jia, W. Huo, Y. Li, T. Zhang, X. Wang, X. Wen, X. Xu, H. Sun, X. Wu, C. Rao, Z. Wei, Z. Zhai, H. Tan, Specialized NADPH diaphorase membrane-related localizations in the brainstem of the pigeons (Columba livia), bioRxiv, DOI 10.1101/663310(2019) 663310.

[27] J.A.L. Clarke, XXII. Further researches on the grey substance of the spinal cord, Philosophical Transactions of the Royal Society of London, 149 (1859) 437–467.

[28] C.S.R. Y, Histologie du systeme nervoux de l’homme et des vertebres, Maloine, 1 (1909) 986.

[29] Y. Li, W. Hou, Y. Jia, C. Rao, Z. Wei, X. Xu, H. Li, F. Li, X. Wang, T. Zhang, J. Sun, H. Tan, Megaloneurite, a giant neurite of vasoactive intestinal peptide and nitric oxide synthase in the aged dog and identification by human sacral spinal cord, bioRxiv, DOI 10.1101/726893(2019) 726893.

[30] P. Bach-y-Rita, Nonsynaptic diffusion neurotransmission (NDN) in the brain, Neurochemistry international, 23 (1993) 297–318.

[31] K. Storer, J. Toh, M.A. Stoodley, N.R. Jones, The central canal of the human spinal cord: a computerised 3-D study, The Journal of Anatomy, 192 (1998) 565–572.

[32] H. Tan, J. He, S. Wang, K. Hirata, Z. Yang, A. Kuraoka, M. Kawabuchi, Age-related NADPH-diaphorase positive bodies in the lumbosacral spinal cord of aged rats, Archives of histology and cytology, 69 (2006) 297–310.

[33] R.B. Leonard, D.H. Cohen, A cytoarchitectonic analysis of the spinal cord of the pigeon (Columba livia), The Journal of comparative neurology, 163 (1975) 159–180.

[34] J. Rosenberg, R. Necker, Fine structural evidence of mechanoreception in spinal lumbosacral accessory lobes of pigeons, Neuroscience letters, 285 (2000) 13–16.

[35] R. Necker, Mechanosensitivity of spinal accessory lobe neurons in the pigeon, Neuroscience letters, 320 (2002) 53–56.

[36] M. Cifuentes, S. Rodríguez, J. Pérez, J.M. Grondona, E.M. Rodríguez, P. Fernández-Llebrez, Decreased cerebrospinal fluid flow through the central canal of the spinal cord of rats immunologically deprived of Reissner’s fibre, Experimental brain research, 98 (1994) 431–440.

[37] T.H. Milhorat, R.W. Johnson, W.D. Johnson, Evidence of CSF flow in rostral direction through central canal of spinal cord in rats, Hydrocephalus, Springer 1991, pp. 207–217.

[38] M. Sakata, K. Yashika, P.H. Hashimoto, Caudal aperture of the central canal at the filum terminale in primates, Kaibogaku Zasshi, 68 (1993) 213–219.

[39] T. Milhorat, S. Nakamura, I. Heger, F. Nobandegani, Ultrastructural evidence of sink function of central canal of spinal cord as demonstrated by clearance of horseradish peroxidase, PROCEEDINGS OF THE ANNUAL MEETING-ELECTRON MICROSCOPY SOCIETY OF AMERICA, SAN FRANCISCO PRESS, 1992, pp. 700–700.

[40] Y. Ikegami, F. Morita, Cerebrospinal fluid contacting neurons and openings of the central canal in rabbits and monkeys--light and electron microscopic observation, Bulletin of the Osaka Medical School, 33 (1987) 1–19.

[41] N. Dale, A. Roberts, O.P. Ottersen, J. Storm-Mathisen, The development of a population of spinal cord neurons and their axonal projections revealed by GABA immunocytochemistry in frog embryos, Proceedings of the Royal Society of London. Series B, Biological sciences, 232 (1987) 205–215.

[42] J.P. Hugnot, R. Franzen, The spinal cord ependymal region: a stem cell niche in the caudal central nervous system, Front Biosci (Landmark Ed), 16 (2011) 1044–1059.

[43] M. Hristov, B. Landzhov, K. Yakimova, Increased NADPH-diaphorase reactivity in the hypothalamic paraventricular nucleus and tanycytes following systemic administration of leptin in rats, Acta Histochem, 121 (2019) 690–694.

[44] C.N. Honda, Visceral and somatic afferent convergence onto neurons near the central canal in the sacral spinal cord of the cat, Journal of neurophysiology, 53 (1985) 1059–1078.

[45] M. Cifuentes, P. Fernández-LLebrez, J. Perez, J. Perez-Figares, E. Rodriguez, Distribution of intraventricularly injected horseradish peroxidase in cerebrospinal fluid compartments of the rat spinal cord, Cell and tissue research, 270 (1992) 485–494.

[46] N.N.Y. Nawar, Observations on the pia-arachnoid of the anterior median fissure and the posterior median sulcus of the fetal spinal cord of the albino mouse, Cells Tissues Organs, 108 (1980) 30–33.

[47] J.A. Foster, P.E. Phelps, Neurons expressing NADPH-diaphorase in the developing human spinal cord, The Journal of comparative neurology, 427 (2000) 417–427.

[48] C. Abbadie, K. Skinner, I. Mitrovic, A.I. Basbaum, Neurons in the dorsal column white matter of the spinal cord: complex neuropil in an unexpected location, Proceedings of the National Academy of Sciences of the United States of America, 96 (1999) 260–265.

[49] E. Jalalvand, B. Robertson, H. Tostivint, P. Löw, P. Wallén, S. Grillner, Cerebrospinal fluid-contacting neurons sense pH changes and motion in the hypothalamus, The Journal of Neuroscience, DOI 10.1523/jneurosci.3359-17.2018(2018).

[50] D.P. Becker, J.A. Wilson, G.W. Watson, The spinal cord central canal: response to experimental hydrocephalus and canal occlusion, Journal of neurosurgery, 36 (1972) 416–424.

[51] B. Vigh, M.J. Manzano e Silva, C.L. Frank, C. Vincze, S.J. Czirok, A. Szabo, A. Lukats, A. Szel, The system of cerebrospinal fluid-contacting neurons. Its supposed role in the nonsynaptic signal transmission of the brain, Histology and histopathology, 19 (2004) 607–628.

[52] K. Chung, W.T. Lee, Vasoactive intestinal polypeptide (VIP) immunoreactivity in the ependymal cells of the rat spinal cord, Neuroscience letters, 95 (1988) 1–6.

[53] C.C. LaMotte, Vasoactive intestinal polypeptide cerebrospinal fluid-contacting neurons of the monkey and cat spinal central canal, The Journal of comparative neurology, 258 (1987) 527–541.

[54] Y. Li, Z. Wei, Y. Jia, W. Hou, Y. Wang, S. Yu, G. Shi, G. Du, H. Tan, Dual forms of aging-related NADPH diaphorase neurodegeneration in the sacral spinal cord of aged non-human primates, bioRxiv, DOI 10.1101/527358(2019) 527358.

[55] S. Rodriguez, S. Hein, R. Yulis, L. Delannoy, I. Siegmund, E. Rodriguez, Reissner’s fiber and the wall of the central canal in the lumbo-sacral region of the bovine spinal cord. Comparative immunocytochemical and ultrastructural study, Cell Tissue Res, 240 (1985) 649–662.

[56] B.T. Hope, G.J. Michael, K.M. Knigge, S.R. Vincent, Neuronal NADPH diaphorase is a nitric oxide synthase, Proceedings of the National Academy of Sciences of the United States of America, 88 (1991) 2811–2814.

[57] T.M. Dawson, D.S. Bredt, M. Fotuhi, P.M. Hwang, S.H. Snyder, Nitric oxide synthase and neuronal NADPH diaphorase are identical in brain and peripheral tissues, Proceedings of the National Academy of Sciences of the United States of America, 88 (1991) 7797–7801.

[58] R. Krstic, D. Nicolas, Presence of Calmodulin and NADPH-Diaphorase Activity but not of Neuronal Nitric Oxide Synthase in the Subcommissural Organ of the Mongolian Gerbil, Cells Tissues Organs, 154 (1995) 232–235.

